# RF Heating of Bipolar Epicardial Implants during MRI at 0.55 T and 1.5T: Effect of Device Length and Termination Conditions

**DOI:** 10.64898/2026.05.26.728047

**Authors:** Bhumi Bhusal, Pia Panravi Sanpitak, Fuchang Jiang, Gregory Webster, Jacob Richardson, Nicole Seiberlich, Laleh Golestanirad

## Abstract

Pediatric patients with epicardial cardiac implantable electronic devices (CIEDs) are frequently excluded from the Magnetic Resonance Imaging (MRI) primarily due to RF heating safety concerns. In this study we evaluate RF heating of two bipolar epicardial leads during MRI at 0.55 T and 1.5 T under different termination conditions. Our findings showed that the mean RF heating was significantly reduced at 0.55 T MRI compared to that at 1.5 T. Similarly, the RF heating at 0.55 T MRI was highest for full system whereas, during MRI at 1.5 T, the RF heating was highest for the capped abandoned lead, showing dependence of RF heating pattern on MRI field strength. While RF heating at both fields surpassed the safety limit, the capped abandoned leads at 1.5 T MRI showed significantly higher RF heating with temperature rise surpassing 50°C in some of the cases. These results highlight the difference in RF heating of bipolar epicardial leads compared to the previously reported findings for monopolar epicardial lead which showed smallest heating for capped abandoned lead at both field strengths. These findings emphasize the necessity of device-specific evaluations at each field-strength to inform clinical decision-making and expand MRI access for this vulnerable population.

## I. Introduction

Congenital heart defects (CHDs) represent the most common form of congenital anomaly in the United States, affecting approximately 1 out of every 100-children born [1]. A substantial proportion of patients with CHDs require cardiac implantable electronic devices (CIEDs), including pacemakers and implantable cardioverter-defibrillators, for rhythm management early in life [2], [3]. While transvenous endocardial lead implantation is standard practice in adults, this approach is frequently unsuitable for pediatric patients due to small vessel diameter [3], [4]. Additionally, for some of the patients, the defects themselves limit the intravenous access to the heart [5]. As a result, epicardial lead implantation—where pacing or sensing leads are surgically attached to the outer myocardial surface—remains the preferred solution for majority of pediatric CIED patients.

Despite the widespread clinical adoption, no commercially available epicardial CIED systems are currently labeled as MR conditional. Consequently, pediatric patients with epicardial implants are routinely excluded from Magnetic Resonance Imaging (MRI), despite MRI being a cornerstone modality for longitudinal evaluation of congenital heart disease and associated comorbidities. The primary safety concern underlying this exclusion is radiofrequency (RF)-induced heating of the implant, particularly in the vicinity of conductive lead electrodes, which may cause permanent tissue damage [6], [7]. In the absence of MRI access, these patients often undergo repeated computed tomography or X-ray examinations, resulting in cumulative exposure to ionizing radiation during a critical period of growth and development of such young patients. A recent study has shown that the pediatric patients with CIEDs are exposed to significantly higher levels of radiation compared to the ones without the implants [8].

While epicardial pacing devices in commercial market exist in both unipolar and bipolar models, the vast majority of pediatric CIED patients undergo bipolar epicardial lead implantation. Even for the same length models, it is possible that the bipolar leads exhibit completely different RF heating characteristics during MRI compared to the unipolar version due to the difference in device geometry.

Recently, MRI systems operating at lower static magnetic field strengths, such as 0.55 T, have gained increasing attention as a potentially safer alternative for imaging patients with implanted medical devices. Compared to conventional 1.5 T or 3 T scanners, low-field MRI systems exhibit reduced magnetic susceptibility related artifacts along with substantially reduced magnetic forces. Additionally, reduced RF frequency at these field strengths leads to reduction in the whole-body specific absorption rate (SAR) for a given applied RF magnetic field (B1), suggesting a reduced RF power deposition for comparable excitation conditions. These characteristics collectively contribute to the perception of improved “implant-friendliness” at low field strengths. However, given the complexity of dependence of implant RF heating during MRI on multiple factors beyond the magnitude of incident electric field, device specific evaluations are required before making any safety assumption during MRI scanning at the low fields [9]–[15]. One important aspect of RF heating of elongated conducting implants is the resonance phenomenon, governed by the antenna behavior of the lead as it couples with the incident RF electric field. This phenomenon may lead to significant amplification of power deposition even with the exposure with lower magnitude electric fields [13], [16], [17].

In this study, we systematically investigate RF-induced heating of bipolar epicardial cardiac implants during MRI at 0.55 T and 1.5 T. For this, we use two bipolar epicardial leads (two different lengths) with different termination conditions: full system (proximal end connected to IPG) and abandoned lead (proximal end capped or uncapped). By comparing heating at low-field and conventional field strengths under matched RF exposure conditions, this work aims to clarify whether and under what circumstances low-field MRI offers a meaningful safety advantage for pediatric patients with bipolar epicardial CIEDs. The findings provide critical insights toward evidence-based MRI safety assessment and may help in making informed clinical decision-making for this vulnerable patient population.

## II. Methods

### A. Devices and Configurations

For this study, we used commercially available bipolar epicardial devices from Medtronic (Medtronic Inc., Minneapolis, Minnesota, USA). We selected two bipolar epicardial leads from Medtronic (models 4968-25 and 4968-35) representing two different lead lengths: 25 cm and 35 cm respectively, which are the most used lead lengths at our institution. For the full system evaluations, the proximal end of the lead was connected to an implantable pulse generator (Medtronic Azure XT DR) whereas for the case of abandoned leads, the lead end was either capped with manufacturer provided insulation cap made of non-conducting plastic material (capped abandoned lead) or left uncapped (uncapped abandoned lead). The device was placed inside an ASTM phantom filled with tissue mimicking gel (conductivity = 0.5 S/m, relative permittivity = 88 measured at 64 MHz frequency). The device was positioned along a particular trajectory on a custom-made rotating disc which was rotated clockwise at an interval of 30 degrees to create 12 different configurations. The measurement setup and starting lead configuration of each lead model has been shown in Fig. 1. The measurements were repeated for all different termination conditions during RF exposure at both 0.55 T and 1.5 T scanners. latex

**Fig. 1.**
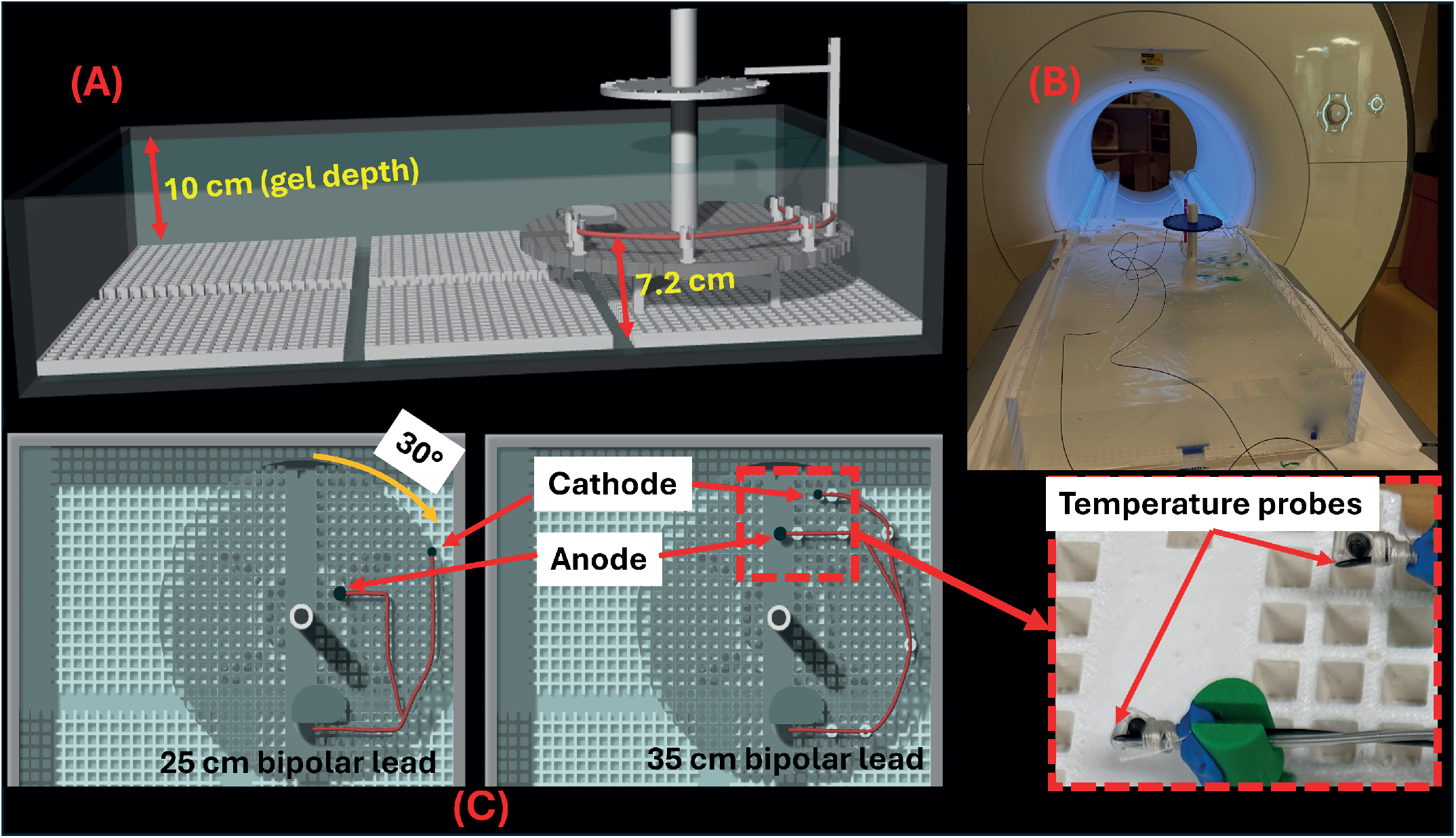
Experimental setup for the MRI induced RF heating measurement experiments. (A) Rectangular ASTM phantom filled with polyacrylic acid gel and rotating platform setup. (B) Phantom setup on the MRI scanner table during measurement. (C) Close view of the starting lead configuration (ID1) for 25 cm (left) and 35 cm (right) bipolar epicardial leads showing temperature probes attached to anode and cathode ends. The rotating disc with the trajectory was rotated in 30^*°*^ intervals to create the rest of the configurations.

### B. RF Exposure and Temperature Measurements

The experimental measurements of MRI induced RF heating were performed at 1.5 T Aera scanner and 0.55 T Free.Max scanner from Siemens (Siemens Healthineers, Erlangen, Germany). At both scanners, we used high-SAR sequences for maximizing the RF heating at the lead-electrodes. For 0.55 T Free.Max scanner, we used a high-SAR T1-saturation pulse sequence with the following scan parameters: acquisition time (TA) = 4:21 min, repetition time (TR) = 671 ms, echo time (TE) = 13 ms, flip angle (FA) = 180°, and 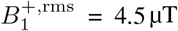. Similarly, for scanning at 1.5 T Aera scanner, we used a high-SAR turbo-spin-echo (TSE) sequence with scanning parameters as follows: acquisition time (TA) = 4:21 min, repetition time (TR) = 836 ms, echo time (TE) = 7.3 ms, flip angle (FA) = 165°, and 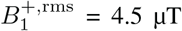. It should be noted that the 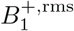 and acquisition time were kept identical for comparison of heating at same level of RF exposure. While sequence parameters were adjusted to maximize RF power/SAR, certain clinical protocols may still reach similar power levels [18].

Fluoroptic temperature probes (OSENSA INC., BC, Canada) with high resolution (0.01°C) were attached to each of the electrodes of the bipolar epicardial leads used in the study. The bipolar lead connected to the temperature probes at both electrodes (cathode and anode) has been depicted in Fig. 1.

## III. RESULTS

The temperature rise measured at the cathode and anode ends of the 25-cm bipolar lead with different termination conditions, during MRI scanning at 0.55 T and 1.5 T scanner have been depicted in the bar plots in Fig. 2. Similarly, the bar plot of temperature increase for different termination condition for a 35-cm bipolar lead has been shown in Fig. 3. The plots indicate that the RF heating at the lead electrode can be strongly dependent on the termination conditions for either of the lead lengths. Interestingly, the RF heating at 0.55 T for both bipolar leads (25 cm and 35 cm length) was highest for the full system (lead connected to IPG) followed by the uncapped abandoned lead and the capped abandoned lead showing the least heating. At 0.55 T, the cathode end which showed higher heating had mean RF heating of 3.44±2.50°C for full system, 1.62±1.21°C for uncapped and 0.68±0.55°C for capped abandoned lead for 25 cm bipolar lead and 4.65±2.86°C for full system, 2.11±1.32°C for uncapped and 1.22±0.89°C for capped abandoned lead for 35 cm bipolar lead. The anode end also showed similar pattern but smaller heating. On the other hand, the heating pattern was completely different for both leads at 1.5 T, where the RF heating was significantly higher for capped abandoned lead followed by the uncapped abandoned lead and the lowest heating observed for the full system. The mean temperature increase at the cathode end at this field strength were 8.86±6.51°C for full system, 9.57±6.76°C for uncapped and 15.14±14.15°C for capped abandoned lead for 25 cm bipolar lead and 3.75±2.69°C for full system, 10.27±5.52°C for uncapped and 34.49±25.01°C for capped abandoned lead for 35 cm bipolar lead.

**Fig. 2.**
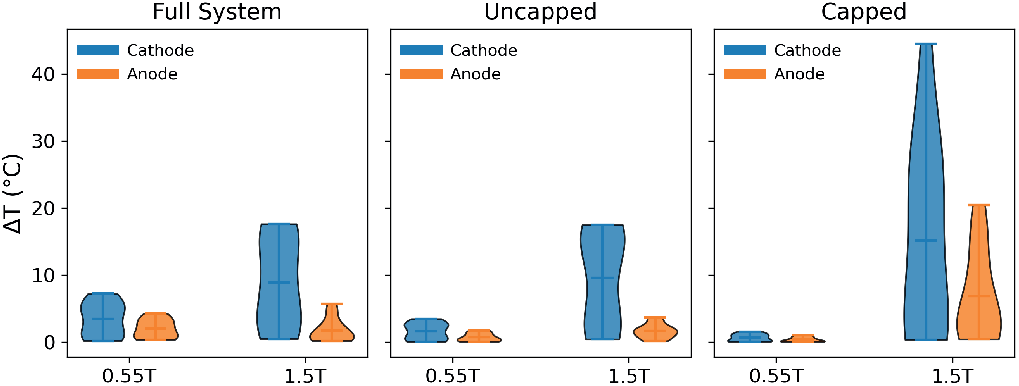
Violin plot showing distribution of temperature increase at the anode and cathode ends of a 25 cm bipolar epicardial lead with different termination conditions during MRI scanning at 0.55 T and 1.5 T.

**Fig. 3.**
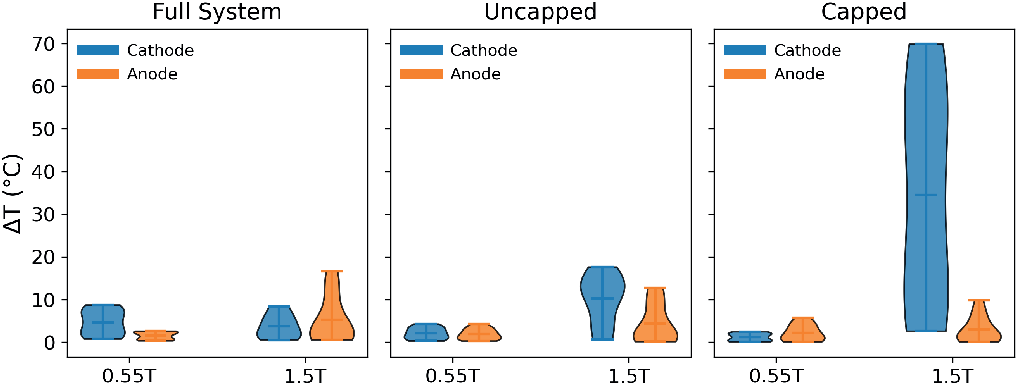
Violin plot showing distribution of temperature increase at the anode and cathode ends of a 35 cm bipolar epicardial lead with different termination conditions during MRI scanning at 0.55 T and 1.5 T.

Fig. 4 represent the comparison of RF heating at the cathode end of the 25 cm and 35 cm bipolar epicardial leads at different termination conditions and field strengths. The plots show similar heating range for 25 cm and 35 cm leads during MRI at 0.55 T. However, at 1.5 T the length effect seems more evident with 25 cm lead showing higher heating for full system and the 35 cm lead showing higher heating for capped abandoned lead. Moreover, the RF heating tends to be substantially smaller at 0.55 T compared to that at 1.5 T (more than 20-fold reduction for the capped abandoned lead, 5-fold reduction for uncapped abandoned lead and up to 2 fold reduction for the full system).

**Fig. 4.**
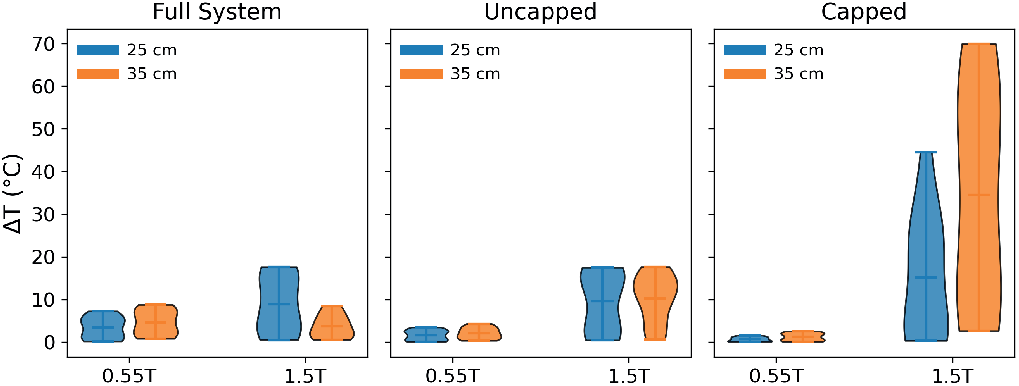
RF heating at the cathode end compared between 25 cm and 35 cm bipolar leads for all termination conditions during MRI at 0.55 T and 1.5 T.

## IV. DISCUSSIONS AND CONCLUSIONS

RF induced heating of elongated conducting implants during MRI is an highly intricate phenomenon, governed by multiple factors such as field strength (RF frequency), implant configurations as well as dielectric properties of surrounding media [11], [12], [14], [19]–[26]. Seemingly, similar implants with different internal geometries or different termination conditions may produce drastically different RF heating behavior even under identical RF exposure in identical scanners. This behavior can be explained by the change in resonance heating behavior determined by antenna characteristics of the implant, which depend on the frequency as well as phase of incident RF fields that couple with the implant lead. Hence, device specific evaluations are of paramount importance while making any decision upon using MRI on patients with such implants.

In this work, we evaluated the RF heating of two commonly used bipolar epicardial leads with length 25 cm and 35 cm to analyze the effect of field strength as well as termination conditions of the leads. Our findings suggest that, the MRI induced RF heating of bipolar-epicardial implants are generally lower at 0.55 T MRI scanner compared to that at 1.5 T MRI. These findings come in agreement with the previous reported studies comparing RF heating of cardiac and neuromodulation implants at these two field strengths [9], [27]. However, this should not be generalized to assume that the lower field MRI is always safe for such patients, as temperature increase far exceeding the risk level were recorded, with some cases having temperature increase approaching 10°C. While, temperature increase below 2°C are considered safe, temperature increase beyond 5°C can cause tissue damage within 10 minutes of RF exposure [28].

Another important finding from this study is that the RF heating for different termination conditions of the epicardial leads is dependent on lead models and field strengths. The findings agree well with an earlier study which reported dependence of the lead-tip RF heatng on the lead lengths as well as termination conditions [29]. For the lower field strength MRI (0.55 T), the RF heating was highest for the full system (lead connected to IPG) and lowest for the capped abandoned lead for both of the bipolar lead models. However, at 1.5 T MRI, the capped abandoned lead produced precariously high heating for both lead lengths, with temperature rise reaching beyond 50°C in some cases. These results with 25 cm bipolar lead show different trend compared to a 25 cm unipolar epicardial lead reported in an earlier study, which showed lowest heating with the capped abandoned lead at both field strengths [27]. These findings emphasize the importance of device-specific RF heating evaluations and dangers of using generalizations in MRI induced RF heating safety. Also, it was observed that the RF heating was generally smaller for the anode branch compared to cathode branch. This is likely be due to placement of lead trajectories where cathode branch was closer to the edge of the phantom, thus exposed to higher magnitude of electric field. More controlled studies will be performed in future to evaluate if there is any difference in heating between the branches when same magnitude electric fields are applied.

While the findings from this study provide important information about RF heating behavior of bipolar epicardial leads at different field strength with varying termination conditions, the results are preliminary and need more detailed evaluation for further generalizations. Our future work will be extending the studies to analyze the RF heating of the implants in heterogeneous patient-specific body models to evaluate the generalizability of the results into realistic patient configurations.

## Acknowledgment

The authors would like to acknowledge the in-kind donation of CIED devices from Medtronic, and the Center for Translational Imaging (CTI) at Northwestern University for the availability of MRI scanners for this study.

